# Optogenetic control of Nodal signaling patterns

**DOI:** 10.1101/2024.04.11.588875

**Authors:** Harold M. McNamara, Bill Z. Jia, Alison Guyer, Vicente J. Parot, Caleb Dobbs, Alexander F. Schier, Adam E. Cohen, Nathan D. Lord

## Abstract

A crucial step in early embryogenesis is the establishment of spatial patterns of signaling activity. Tools to perturb morphogen signals with high resolution in space and time can help reveal how embryonic cells decode these signals to make appropriate fate decisions. Here, we present new optogenetic reagents and an experimental pipeline for creating designer Nodal signaling patterns in live zebrafish embryos. Nodal receptors were fused to the light-sensitive heterodimerizing pair Cry2/CIB1N, and the Type II receptor was sequestered to the cytosol. The improved optoNodal2 reagents eliminate dark activity and improve response kinetics, without sacrificing dynamic range. We adapted an ultra-widefield microscopy platform for parallel light patterning in up to 36 embryos and demonstrated precise spatial control over Nodal signaling activity and downstream gene expression. Patterned Nodal activation drove precisely controlled internalization of endodermal precursors. Further, we used patterned illumination to generate synthetic signaling patterns in Nodal signaling mutants, rescuing several characteristic developmental defects. This study establishes an experimental toolkit for systematic exploration of Nodal signaling patterns in live embryos.

## Introduction

Embryos often transmit instructions to their cells using concentration-dependent signaling cues called morphogens. Spatial patterns of morphogen concentration convey positional information to cells, activating position-appropriate developmental programs^1–7^. Precisely how cells extract this information from morphogen distributions remains an open question^8,9^. In the classical model, each cell autonomously measures its local signal concentration and selects the appropriate fate in response^4,5,8^. However, it has become clear that cells often go beyond simple concentration sensing and instead respond to more complex features of morphogen patterns. For example, cells can pool information via secreted signals to sense signaling domain size in ‘community effects’^10,11^ or modify their decisions based on geometric features of their community structure^12^. Morphogen dynamics can also carry information; cells can respond differently depending on exposure timing and duration^13–21^ or whether signaling accumulates abruptly or slowly^22,23^. Morphogen responses can also be probabilistic, such that a cell’s signaling history determines its fate only in a statistical sense. Indeed, patterning of the zebrafish endoderm and neural tube are initially noisy, only to be refined by downstream processes^24–26^.

Testing quantitative theories of how morphogens organize development requires the ability to systematically manipulate spatial and temporal patterns of signaling activity. Traditional methods can achieve coarse perturbations. For example, genetic knockouts can remove or expand morphogen domains^27,28^, and microinjections or transplants can introduce point sources of morphogen cues^29,30^. However, the lack of precise spatial and temporal control makes it difficult to explicitly test patterning models. Ideally, an investigator could design and create arbitrary morphogen signaling patterns—in time and space— to rigorously test specific hypotheses.

Optogenetic tools have emerged as a promising strategy for agile and precise control over developmental gene expression^31–33^ and signaling^34–36^. In an approach pioneered in receptor tyrosine kinase signaling^37^, active signaling complexes are assembled by tagging components with protein domains that dimerize in response to light. By rewiring signaling pathways to respond to light, one can, in effect, convert photons into morphogens. Modern optical techniques, in turn, allow for light patterning with sub-millisecond time resolution and subcellular spatial resolution^38–40^. In principle, these tools unlock a level of control over developmental signaling that cannot be achieved with traditional manipulations.

In developmental biology, optogenetic strategies have been applied most extensively to investigate terminal patterning via the Ras/ERK signaling pathway in the early Drosophila embryo^14,41–43^. These approaches have now been applied to several morphogen pathways^44,45,46^ as well as to vertebrate embryos^47,21,48^; however, practical challenges have prevented widespread adoption. First, optogenetic reagents often suffer from limited dynamic range. To mimic developmental signaling patterns with light, an optogenetic reagent must switch from negligible background activity in the dark to light-activated signaling levels approaching peak endogenous responses. Second, common strategies for spatial light control have limited throughput and flexibility. Systematic dissection of morphogen signaling mechanisms requires a means to deliver precise patterns of light to large numbers of live embryos as they grow and change shape.

Nodal is a TGFβ family morphogen that organizes mesendodermal patterning in vertebrate embryos^49–52^. Nodal ligands exert their effects by assembling complexes of Type I and Type II cell surface receptors and an EGF-CFC family cofactor^49,53–55^. Ligand-induced proximity between the receptors leads the constitutively-active Type II receptor to phosphorylate and activate the Type I receptor, which then phosphorylates the transcription factor Smad2^56^. Once active, pSmad2 translocates to the nucleus and, in concert with other transcriptional cofactors, induces the expression of Nodal target genes^57,58^. In zebrafish, the Nodal ligands Cyclops and Squint are produced at the embryonic margin^59–62^. Cyclops and Squint dimerize with the ubiquitously expressed Nodal ligand Vg1 prior to secretion to form active heterodimeric ligands^63–65^. Diffusion of these ligands from the margin generates a vegetal to animal concentration gradient that instructs germ layer fate selection^28,30,50^; higher Nodal exposure directs cells to endodermal fates, while lower levels direct cells to mesodermal fates^15,50,59,66–68^. Recent work also suggests that the Nodal signaling gradient establishes a gradient of cell motility and adhesiveness that is important for ordered cell internalization at the onset of gastrulation^69,70^.

Nodal was the first developmental signal to be made optogenetically tractable in zebrafish through fusion of the Type I and Type II receptors *acvr1b* and *acvr2b* to the photo-associating LOV domain of aureochrome1 of the alga *Vaucheria frigida*^21,71^. Under blue light illumination, dimerization of the LOV domains brings the receptors together and initiates signaling. While these first-generation ‘optoNodal’ reagents enabled temporal control of Nodal target gene expression, spatial patterning of Nodal signaling with light has not yet been reported. Further, LOV domains often exhibit slow dissociation kinetics^72^ which may limit the temporal resolution with which signals can be controlled, and may also contribute to problematic dark activity. Achieving biologically relevant spatial patterning places more stringent technical requirements on both optogenetic reagents and optical instrumentation than does temporal patterning.

Here we report an experimental pipeline for optogenetic patterning of Nodal signaling with improved dynamic range, as well as higher temporal resolution, spatial resolution, and throughput. We develop improved optoNodal reagents (hereafter optoNodal2) with enhanced dynamic range by fusing Nodal receptors to the light-sensitive heterodimerizing pair Cry2/CIB1N, and by further sequestering the Type II receptor to the cytosol. We use a custom ultra-widefield patterned illumination approach^73^ for spatial patterning and live imaging of up to 36 zebrafish embryos in parallel. We demonstrate flexible patterning of Nodal signaling activity and target gene expression in zebrafish embryos. We further demonstrate spatial control over cell internalization movements during gastrulation, and rescue of several development defects in Nodal signaling mutants. Our platform lays the foundation to systematically dissect the spatial logic of Nodal signaling and demonstrates a generalizable approach to high-throughput optogenetic control over morphogen signals in the zebrafish embryo.

## Results

### Development of new optoNodal reagents with enhanced kinetics and dynamic range

An ideal optogenetic reagent would evoke strong signaling in response to light and no signaling in the dark. In practice, many photo-associating domains exhibit some affinity in the dark, leading to unwanted background activity. The original, LOV-based, optoNodal reagents were highly active in the light, as they were able to induce robust expression of ‘high-threshold’ Nodal expression targets such as *gsc* and *sox32*^21^. However, we noticed problematic levels of dark activity even when expressed at low doses of mRNA; wild-type zebrafish embryos injected with LOV optoNodal mRNAs and raised in the dark exhibited measurable Nodal signaling activity as visualized by pSmad2 immunostaining as well as severe phenotypes at 24 hpf, consistent with hyperactive Nodal signaling (Fig. S1a).

We set out to design improved optoNodal receptors (Fig. 1A,B). Inspired by a recent study on optogenetic TGFβ receptors^74^, we reasoned that dark activity could be reduced by introducing two modifications. First, we replaced the LOV-based photo-associating domains with photo-associating domains from *Arabidopsis* Cry2 and Cib1, which have previously been used to engineer light-driven dimerization events with rapid association (∼seconds) and dissociation (∼minutes)^75^. Second, we removed the myristoylation motif from the constitutive Type II receptor so it became cytosolic in the dark. We hypothesized that this change would decrease the effective concentration at the membrane in the dark, reducing the propensity for spurious, light-independent interactions. Indeed, we found that dark activity is greatly reduced over a wide range of mRNA dosages for the redesigned receptors. Embryos injected with up to 30 pg of mRNA coding for each receptor appear phenotypically normal at 24 hpf when grown in the dark (Fig. S1C,D).

**Fig. 1.**
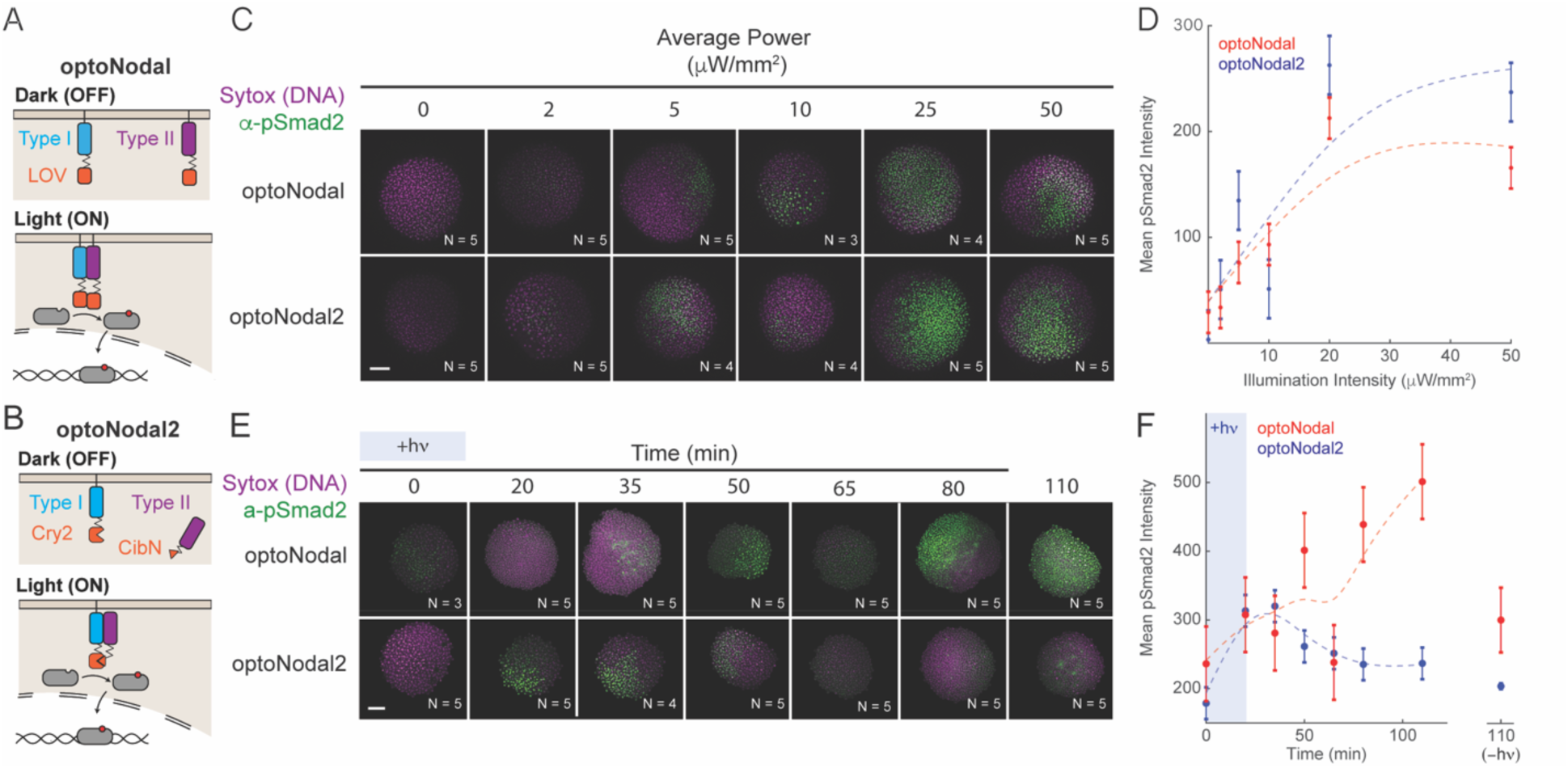
Improved optoNodal2 reagents based on Cry2-Cib1N heterodimerization. (A) Schematic of previously developed LOV-based optoNodal reagents^21^. Type I and Type II receptors are tethered to the membrane via a myristoylation motif (top). Blue light induces homodimerization between LOV domains, activating Nodal signaling (bottom). (B) Schematic of OptoNodal2 reagents. The myristoylation motif is removed from the Type II receptor, localizing it to the cytoplasm (top). Blue light induces heterodimerization of Cry2 and Cib1N, activating Nodal signaling (bottom). (C) Blue light intensity responses for optoNodal (top row) and optoNodal2 (bottom row) reagents. Embryos injected with indicated reagents were illuminated for 1 hour with 470 nm light with the indicated intensity. Nodal signaling was measured by α-pSmad2 immunostaining (green). Images are maximum intensity projections of representative embryos. Scale bar 100 μm. (D) Quantification of Nodal signaling activity from panel C. α-pSmad2 staining intensity was extracted from segmented nuclei in optoNodal (red) and optoNodal2 (blue) treatment groups; each point represents the average nuclear staining intensity from replicate embryos. Number of replicate embryos for each condition are indicated in the relevant images in panel C. Error bars denote the standard error of the mean. Dashed curves depict cubic smoothing spline interpolations. (E) Measurement of response kinetics for optoNodal (top row) and optoNodal2 (bottom row) reagents. Embryos injected with indicated reagents were illuminated for 20 minutes with 470 nm light. Nodal signaling was measured by α-pSmad2 immunostaining (green). Images are maximum intensity projections of representative embryos. (F) Quantification of Nodal signaling activity from panel E. α-pSmad2 staining intensity was extracted from segmented nuclei in optoNodal (red) and optoNodal2 (blue) treatment groups; each point represents the average nuclear staining intensity from replicate embryos. Number of replicate embryos for each condition are indicated in the corresponding images in panel E. Error bars denote the standard error of the mean. Dashed curves depict cubic smoothing spline interpolations. Background intensity of unilluminated embryos at the 110 minute timepoint are included (-hν) to indicate baseline levels of signaling activity.

We next compared the inducibility and kinetics of optoNodal2 relative to previously reported optoNodal. To test the illumination responses, we injected equal amounts of mRNA encoding each set of reagents into mutant embryos lacking endogenous Nodal signaling (M*vg1* mutants) and exposed the embryos to 1 hour of blue light illumination with varying intensity using an open-source LED plate^39^. Both sets of receptors induced Smad2 phosphorylation over a similar range of powers (saturating near 20 μW/mm^2^, Fig. 1. C and D). Notably, the optoNodal2 receptors exhibit equivalent potency without the drawback of detrimental dark activity (Fig. S1A, B). To measure dynamic responses, we exposed M*vg1* embryos expressing the two sets of receptors to a 20-minute impulse of saturating light intensity (20 μW/mm^2^) and stained for pSmad2 at several timepoints following stimulation. The optoNodal2 reagents exhibited rapid kinetic responses; pSmad2 levels reached maximal intensity approximately 35 minutes after stimulation and returned to baseline approximately 50 minutes later. By contrast, signaling in the optoNodal reagents continued to accumulate for at least 90 minutes after cessation of illumination. We confirmed this observation by repeating the dynamic response measurements in an independent mutant background lacking Nodal signaling^53^ (MZ*oep*, Fig. S2). Thus, the optoNodal2 reagents improved the dynamic range and response kinetics over the original optoNodal design without sacrificing potency of light-driven Nodal pathway activation.

### A platform for high-throughput spatial patterning of Nodal signaling activity

Optogenetic tools in developmental biology promise the ability to test spatial and temporal patterns of signaling activity on demand. Recent studies have described spatial modulation of developmental signaling using microscope-coupled digital micromirror devices (DMDs)^42^, well as laser scanning over geometrically-defined regions of interest (ROIs)^47^, and LED illumination with static photomasks^40^. These approaches have limited throughput and flexibility: most DMD-equipped and laser-scanning microscopes can only address a single embryo at a time, and static photomasks require long turnaround times to design and test new patterns. The ability to flexibly pattern signaling in multiple embryos in parallel would open the possibility of systematically exploring how geometric pattern features guide developmental outcomes.

To achieve this goal, we adapted an ultra-widefield microscope system that has been applied to large-area optogenetic manipulation of mouse brain slices^73^ and to study the early electrophysiology of developing zebrafish hearts^76^ (Fig. S4). The microscope leverages a 4x macro objective lens and DMD projector to address a ∼15 mm^2^ area. The system can project light patterns over 8 zebrafish embryos in a single field of view with close-to single-cell resolution. We outfitted the microscope with a scanning stage, multi-color LED illuminator and motorized filter wheel to enable simultaneous multi-channel fluorescence imaging and scanning over multiple fields of view. Further, we built a custom microscope interface in MATLAB that enables custom scripting of each microscope component, paving the way for complex acquisitions that incorporate position scanning, imaging, and spatial light patterning. To mount zebrafish embryos for patterning, we 3D-printed embryo mounts that allow blastula and gastrula stage embryos to be arranged in a regular array (Fig. S3). Embryos mounted in this way remain still enough for precise light delivery, and they develop normally over 24 hours (Supplemental Movie S1).

To demonstrate the spatial patterning capability of this platform, we projected light patterns—a spot, line or bullseye (Fig. 2C)— onto sphere-stage zebrafish embryos mounted in a regular array. Precise Nodal signaling patterns, as read out by pSmad2 immunostaining, could be generated for each pattern with a 20 minute stimulation (Fig. 2D). Application of patterns for longer times (45 minutes) induced spatially patterned gene expression of both a gene in the Nodal regulatory pathway (*lefty2*, Fig 2F) and of a Nodal target gene encoding axial mesodermal fate (*flh*, Fig. 2E). Collectively, these results demonstrate that the new optoNodal2 reagents, coupled with an ultra-widefield patterning platform, enable spatial and temporal patterning of Nodal signaling activity and Nodal-dependent gene expression.

**Fig. 2.**
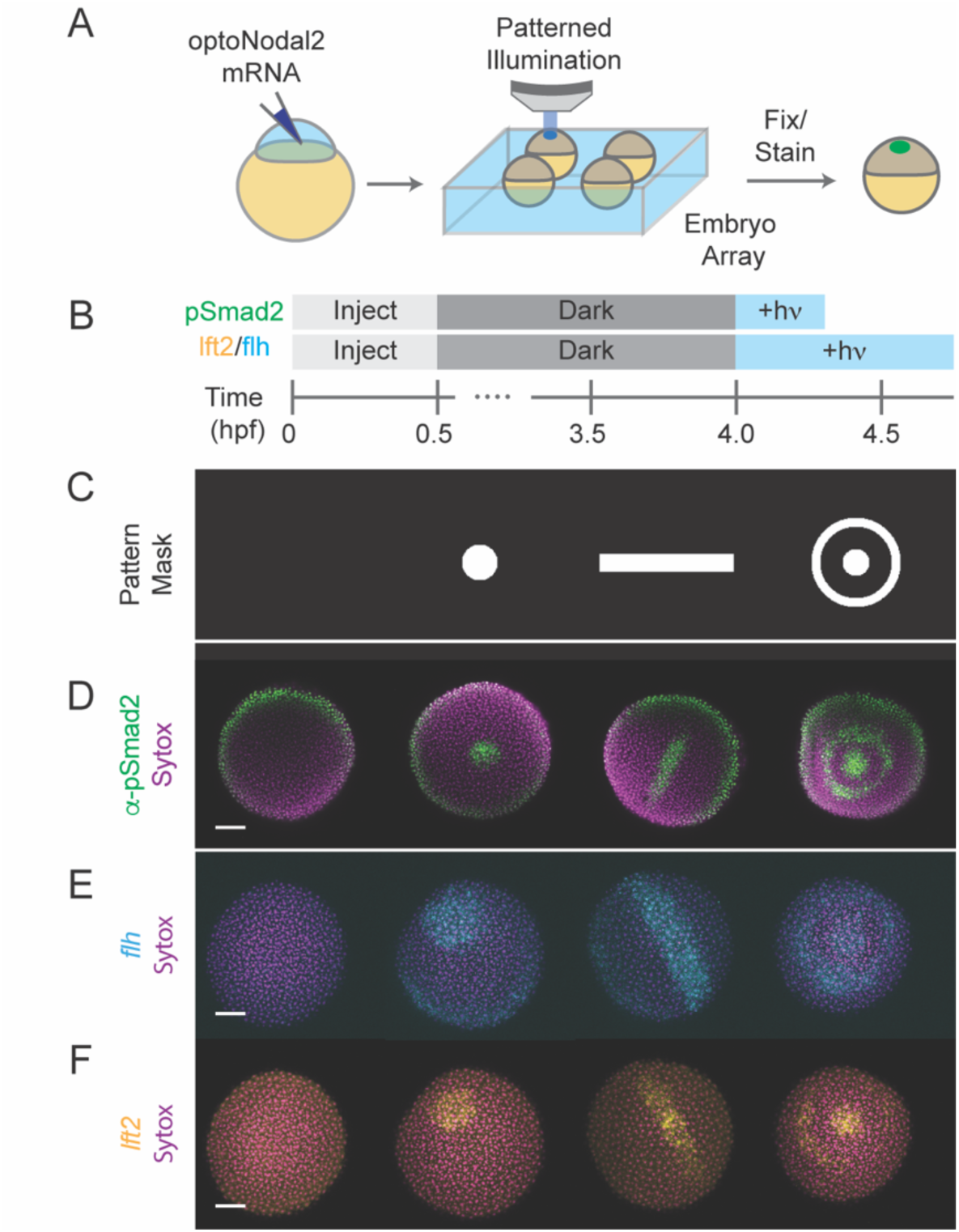
Platform for spatial and temporal patterning of Nodal signaling activity. (A) Schematic of patterning experiment. One-cell embryos were injected with mRNA encoding optoNodal2 receptors. At sphere stage, embryos were mounted in custom array mounts compatible with an upright microscope. Spatial patterns of light were generated using an ultra-widefield microscope incorporating DMD-based digital projector (Fig. S4). (B) Experimental timeline. Embryos were injected with optoNodal2 mRNAs at the 1-cell stage. Embryos were kept in the dark until 4 hpf. Embryos stained for pSmad2 (panel D) were illuminated from 4-4.3 hpf, while embryos stained for *lft2* or *flh* expression (panels E and F) were illuminated from 4-4.75 hpf. All embryos were fixed immediately following light treatment. (C-F) Demonstration of spatial patterning of Nodal signaling activity and target gene expression. (C) DMD pattern masks used for spatial patterning. (D) α-pSmad2 immunostaining (green) demonstrating spatial patterning of signaling activity. (E) Spatial patterning of *flh* gene expression (cyan). (F) Spatial patterning of *lft2* gene expression (yellow). Embryos were double stained for *lft2* and *flh*; each column of images in panels E and F depict the same embryo imaged in different channels. All images in panels D-F are maximum intensity projections derived from confocal images of a representative embryo. All scale bars 100 μm.

### Optogenetic patterning of endodermal cell specification and internalization

We next sought to initiate more complex developmental programs using patterned Nodal stimulation. In zebrafish, endodermal cells are specified by high levels of Nodal signaling within two cell tiers of the margin, after which they internalize via autonomous ingression at the onset of gastrulation^70,77^. We therefore reasoned that optogenetic stimulation targeted to the margin could initiate endodermal specification (i.e. *sox32* expression) and internalization movements in the absence of endogenous Nodal signaling. To test this hypothesis, we injected RNA encoding our optoNodal2 receptors into MZ*oep* mutants and stimulated the margin with targeted illumination from 3.5 hpf (just prior to Nodal signaling onset) until 6 hpf (early gastrulation) (Fig. 3A). To visualize specification, internalization and dispersal of endodermal cells, we harvested stimulated and dark-control embryos at 4 hpf, 6 hpf and 9 hpf, and stained for *sox32* mRNA.

**Fig. 3.**
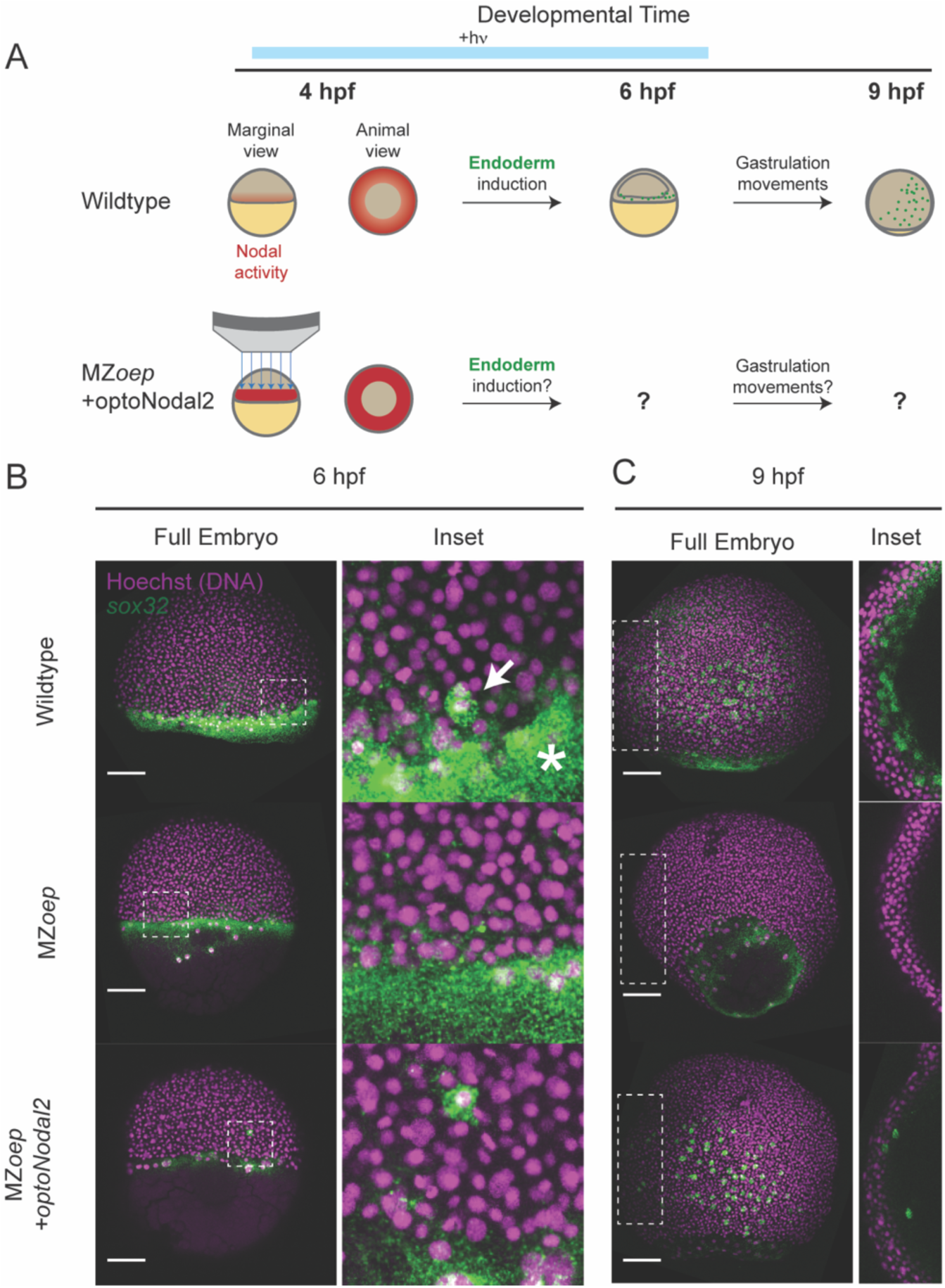
Rescue of endoderm precursors and internalization movements. (A) Endoderm rescue experiment. In wild-type embryos, Nodal signaling near the margin turns on the master endoderm transcription factor *sox32* at 4 hpf. By 6 hpf, *sox32*^+^ endodermal precursors have internalized, and by 9 hpf they have spread over the yolk via random walk movements. MZ*eop* mutants lack Nodal signaling and do not specify endoderm. We rescued *sox32* expression and downstream cell movements in MZ*oep* embryos by targeted optoNodal2 stimulation at the margin from 3.75-6.25 hpf (indicated by blue bar). (B) Rescue of *sox32* expression at 6 hpf expression with optoNodal2 stimulation*. Sox32*^+^ cells were visualized by HCR in wild-type (top row), MZ*oep* (middle row) and optoNodal2-stimulated MZ*oep* embryos (bottom row). Insets and white arrow highlight localization of *sox32^+^* cells at the embryonic margin. Asterisk highlights Nodal-independent *sox32* expression in the extraembryonic yolk syncytial layer. (C) Rescue of cell internalization movements with optoNodal2 stimulation. *Sox32*^+^ cells were visualized by HCR at 9 hpf in wild-type (top row), MZ*oep* (middle row) and optoNodal2-stimulated MZoep (bottom row). Insets depict maximum intensity projections of middle confocal slices to visualize the hypoblast cell layer. *Sox32*^+^ cells reside in the hypoblast in wild-type and optoNodal2-treated embryos at 9 hpf. All scale bars 100 μm.

Confocal imaging of patterned Mz*oep* embryos revealed a salt-and-pepper pattern of *sox32* induction at the margin at 6 hpf, consistent with its expression pattern in wildtype embryos (cf. Fig. 3B, top and bottom rows). Further, we found that at 9 hpf *sox32*^+^ cells in illuminated Mz*oep* embryos had migrated animally and spread over the yolk, again mimicking the normal distribution of endodermal precursors (Fig. 3C). Importantly, individual confocal sections reveal that the induced *sox32*^+^ cells reside in the hypoblast, consistent with them executing internalization movements at gastrulation (Fig. 3C, right column). In unilluminated MZ*oep* mutants, by contrast, *sox32^+^* cells were absent at all observed stages. Collectively, these results demonstrate that we can rescue specification of endodermal precursors and gastrulation-associated internalization movements using targeted optogenetic stimulation.

### Replacement of endogeneous Nodal signaling with patterned illumination

We next tested whether our patterning platform could be used to induce formation of more complex Nodal-dependent tissues. An attractive application of developmental optogenetics is to test which features of morphogen signals are required for downstream development. For example, a recent study in *Drosophila* demonstrated the ability to rescue the development of a lethal patterning mutant using surprisingly simple, optogenetically-evoked spatial patterns of ERK signaling^42^. The capacity to pattern many embryos simultaneously could extend this approach to systematic investigation of how features of spatial patterns encode developmental phenotypes. We therefore tested the ability of a family of stimulation patterns with a range of intensities and spatial extents to rescue the development of MZ*oep* mutants (Fig. 4A,B). We injected MZ*oep* mutants with RNA encoding optoNodal2 reagents and arrayed 36 embryos in our embryo mounts with animal pole facing the microscope objective. We illuminated each embryo with a ‘ring’ pattern that covered the Nodal signaling domain around the embryo margin (Fig. 4C). Pattern characteristics were varied along each dimension of the array; the ring width was varied along one axis (75 μm or 150 μm) and illumination intensity was varied along the other (40 μW/mm^2^, 20 μW/mm^2^ or 10 μW/mm^2^ average intensity (Fig. 4D). Embryos were stimulated from just before the normal onset of Nodal signaling (3.5 hpf) until the onset of gastrulation (6 hpf) to mimic the physiological duration of Nodal signaling. Embryos were collected and raised until 26 hpf in the dark, at which point they were imaged for gross phenotypes.

**Fig. 4.**
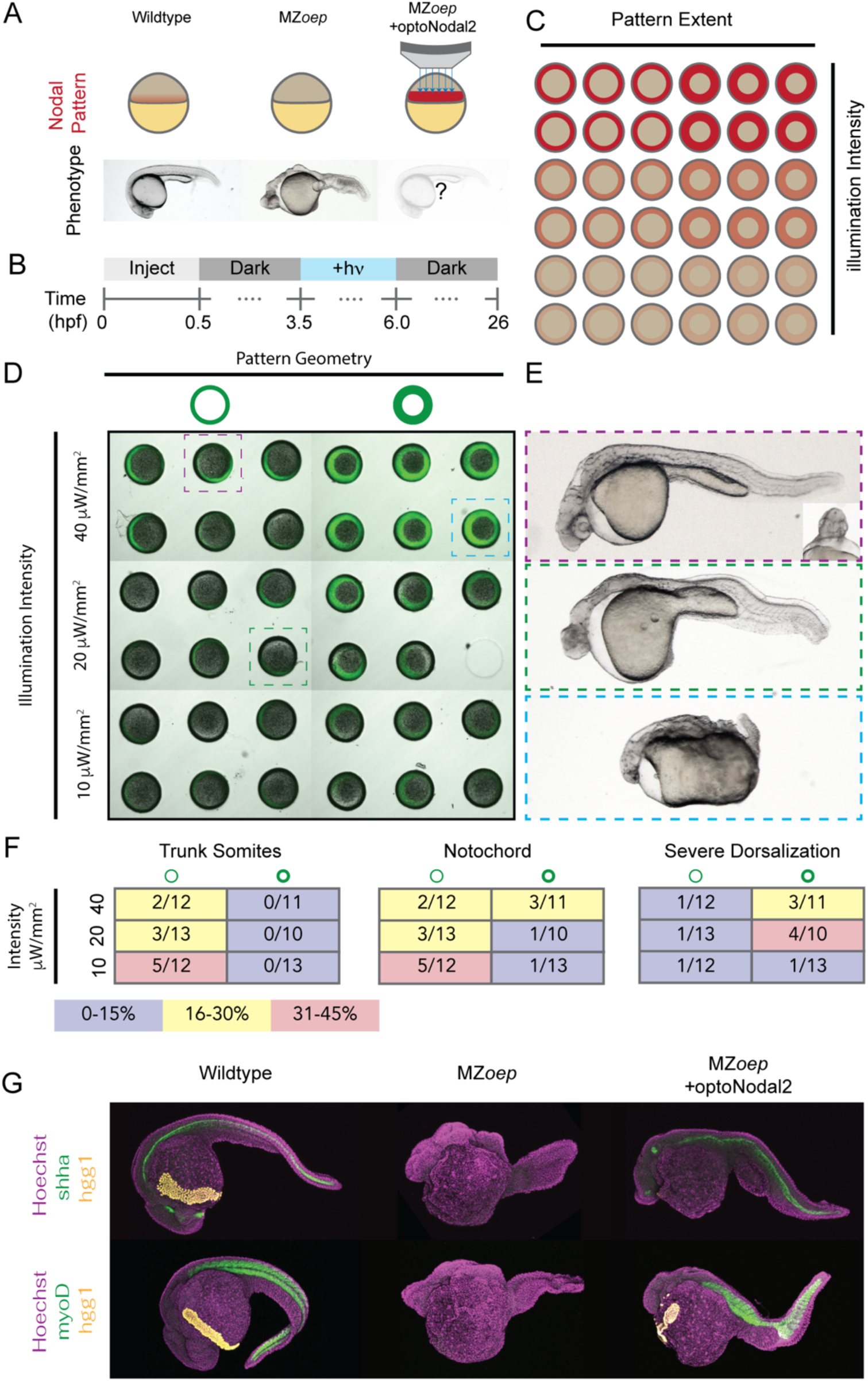
Optogenetic rescue of Nodal signaling mutant phenotypes. (A) Experimental overview. The absence of Nodal signaling in MZ*oep* mutants (middle) results in loss of nearly all mesendodermal tissues. We injected optoNodal2 mRNA into 1-cell stage MZ*oep* embryos and replaced endogenous Nodal signaling with patterned optogenetic stimulation (bottom) (B) Experimental timeline. Illumination patterns were applied from 3.5-6 hpf. Embryos were imaged or fixed at 26 hpf.. (C) Schematic of arrayed layout of Nodal patterns. Optogenetic pattern characteristics were varied along each axis of the embryo array; pattern geometry was varied left-to-right, and pattern intensity was varied top-to-bottom. (D) Visualization of stimulation patterns. Applied patterns (green) were visualized by projecting pattern masks with 560 nm illumination and observing fluorescence from a co-injected mCherry mRNA. Each combination of pattern geometry and intensity was tested in 5 or 6 replicate embryos in the depicted experiment. The 26 hpf phenotypes of boxed embryos are highlighted in panel C. (E) Example rescue phenotypes. Example of a strong rescue phenotype (top), exhibiting notochord, trunk somites and partial rescue of cyclopia. Weaker intensity stimulation (middle) resulted in weaker rescue, with incomplete specification of trunk somites and notochord. Combining high intensity and large-area stimulation led to phenotypes reminiscent of Nodal gain-of-function (bottom, e.g. *lefty1;lefty2* double mutants). (F) Quantification of rescue phenotype frequencies for trunk somites (left), notochord (middle) and severe dorsalization (right). Phenotypes were assessed by visual inspection of transmitted light images. (G) Visualization of marker gene expression for Nodal-dependent tissues. Top row: expression of notochord (*shha*, green) and hatching gland (*hgg1*, yellow) markers. Bottom row: expression of somite (*myoD*, green) and hatching gland (*hgg1,* yellow) markers.

Our treatments elicited different Nodal phenotypes, ranging from a typical MZ*oep* phenotype to rescue of complex structures to phenotypes consistent with Nodal gain of function (e.g. *lefty1;lefty2* double mutants^27^). The frequency of these phenotypes correlated with the characteristics of the applied patterns (Fig. 4F). For example, patterns with lower intensity resulted in higher frequencies of MZ*oep*-like phenotypes (Fig. 4E, middle), whereas thick, intense rings of illumination generated Nodal gain-of-function phenotypes (Fig. 4E, bottom). Complex structures were most often rescued with narrow, low-intensity rings of Nodal activation. In the best examples of rescue, we observe rescue of mesodermally-derived structures such as prechordal plate, notochord and trunk somites (Fig. 4E, top). We confirmed the presence of these tissues in rescued 24 hpf embryos by staining for expression of marker genes for notochord (*shha*), trunk muscle (*myoD*), and hatching glad (*hgg1*), a derivative of the prechordal plate (Fig. 4G). In some embryos, beating heart tissue was observed at the embryonic midline (Supplemental Movie. S2). Finally, with low frequency (2/71 embryos in two replicate experiments) we observe partial rescue of cyclopia, another hallmark of Nodal loss of function mutants (Fig. 4E top, inset). Collectively, these results show that development of complex tissues can be initiated by patterned optogenetic activation of Nodal signaling.

## Discussion

Here, we report the design and application of new optoNodal reagents with reduced dark activity and improved response kinetics. We combine these optoNodal2 reagents with a versatile ultra-widefield optical patterning platform to exert precise control over Nodal signaling activity in time and space, across multiple embryos in parallel. Our pipeline enables optogenetic patterning and subsequent fluorescence imaging of live zebrafish embryos with a substantial improvement in throughput over standard approaches in developmental optogenetics. We demonstrate spatial control of Nodal signaling activity and target gene expression, patterning of endodermal progenitor specification and internalization movements, and phenotypic rescue of Nodal signaling mutants. To our knowledge, the MZ*oep* rescue represents the first application of patterned optogenetics to rescue a mutant phenotype in a vertebrate embryo.

We believe that these improvements will enable optogenetic investigation of new questions requiring stringent spatiotemporal control over Nodal signaling. Indeed, an accompanying study applies our optogenetic system to reveal how Nodal signaling dynamics control convergence and extension movements of the zebrafish mesoderm^78^.

A gradient of Nodal signaling has long been recognized to orchestrate mesendodermal patterning in vertebrate embryos. At first glance, experiments from zebrafish^15,50,59,66,68^ make a compelling case for a concentration threshold-like model; high, medium and low concentrations of Nodal correlate with spatially-ordered populations of endoderm, mesoderm and ectoderm, respectively. However, several experimental observations suggest a more complex picture. The duration of Nodal exposure^13,15^, timing of signal onset and cessation^21^, speed of Nodal spread through space^79^, and kinetics of target gene transcript accumulation^58^ have all been shown to influence fate selection. Complicating matters further, a recent study suggested that Nodal-mediated endoderm fate selection is probabilistic^26^. Our recent observations^28^ suggest that a surprising degree of patterning can be achieved even without a stable gradient. In zygotic *oep* mutants— a background that successfully specifies somites and notochord^68^— the Nodal gradient is transformed into a wave of signaling activity that propagates from the margin toward the animal pole^28^. We do not yet know what constraints the Nodal pattern needs to satisfy or how spatial or temporal features of Nodal signaling allow the embryo to meet them. We anticipate that the approaches we present here will prove useful for answering these questions by enabling Nodal pattern features and dynamics to be manipulated precisely.

Modeling efforts have aimed to explain how diffusion and capture give rise to morphogen profiles^30,80–84^, or how cells transform continuously varying concentrations into discrete fate choices^5^. An enduring challenge with these efforts is that a lack of variation in observed morphogen profiles leaves the models underdetermined. To rigorously constrain quantitative models, we need access to rich libraries of morphogen signaling profiles. For example, models would benefit from datasets that systematically varied signaling gradient range, shape, rate of change, and orientation. In the rare cases where such manipulation is possible surprising outcomes are common. For example, flattened Bicoid gradients performed remarkably well in patterning target gene expression^85^, despite the fact that traditional concentration-centric models predict marked shifts in expression domain boundaries. By making it possible to generate libraries of complex signaling patterns in dozens of embryos simultaneously, we believe that the tools presented here will facilitate rigorous testing of quantitative models.

## Materials and Methods

### Zebrafish husbandry

Zebrafish were raised and maintained according to standard practices^86^. Briefly, embryos were grown in embryo medium (250 mg/L Instant Ocean salt in distilled or reverse osmosis-purified water, adjusted to pH 7.0 with NaHCO3) supplemented with 1 mg/mL methylene blue. Wild-type breeding stocks were the result of TL x AB crosses (TL and AB stocks were obtained from ZIRC). *Vg1* and *oep* mutant fish were propagated as previously described^53,63^. Staging was performed using a combination of time measurement (i.e. time elapsed since fertilization) and morphological examination as compared to a standard staging series^87^. M*vg1* mutant embryos were obtained by mating *vg1^-/-^*females to TLAB wild-type males. MZ*oep* mutant embryos were obtained by incrossing *oep*^-/-^ adult fish. All animal experiments were performed under the supervision of the University of Pittsburgh IACUC (protocol ID 23124380).

### mRNA synthesis and embryo microinjection

Coding sequences for mRNAs used in this study (*myr-acvr1b-cry2*, *acvr2b-cibn*, *myr-acvr1b-LOV*, *myr-acvr2b-LOV*) were cloned into pCS2+ vectors. Briefly, *myr*-*acvr1b-lov* and *myr*-*acvr2b-lov* transcription templates obtained as a gift from the Heisenberg Lab^21^, and Cry2 and Cib1N coding sequences were obtained from Addgene (accession numbers 26866 and 26867, respectively). We replaced the LOV domain sequences in the Myr-Avr1b-LOV and Myr-Acvr2b-LOV coding sequences with Cry2 and Cib1N, respectively, using Gibson assembly cloning. To transcribe mRNA, plasmid templates were linearized with NotI and purified using Monarch PCR purification kits (New England Biolabs). The purified templates were transcribed using mMESSAGE mMACHINE Sp6 (Thermo-Fisher Scientific) kits. mRNAs were purified using the Monarch RNA Cleanup Kit (New England BioLabs) and eluted in RNAse-free water. All kits were used according to manufacturer’s specifications. Plasmids encoding optoNodal2 reagents were deposited with Addgene under accession numbers 161715 and 161720.

mRNA microinjections were carried out using Drummond Nanoject III injector instruments. Injections were performed directly into the blastomere of 1-cell stage dechorionated embryos. Injections were typically 2.0 nL in volume, and embryos raised in agarose-coated 6-well dishes in embryo medium supplemented with methylene blue following injection. All embryos injected with optogenetic reagents were kept in aluminum foil-wrapped plates for all timepoints after 2.0 hpf.

### OptoNodal Receptor light intensity and impulse response measurements

All intensity response and kinetic response measurements from Fig. 1 were obtained using the open-source optoPlate-96 instrument^39^ (dual blue LED configuration). Our instrument was fabricated by the machine shop at the University of Pittsburgh Department of Cell Biology. Power calibration for each LED was performed using a ThorLabs PM100d power meter and a custom MATLAB analysis script.

Experiments were designed—with intensity correction factors applied— and transferred to the optoPlate Arduino processor using the OptoConfig software package^88^. For the light intensity response series (Fig. 1 C,D), we injected 15 pg of either Cry-Cib or LOV optoNodal reagents into M*vg1* embryos at the 1-cell stage. A 1-hour light treatment (average powers of 1,5,10,25 and 50 μW/mm^2^) was initiated at sphere stage. These light pulses consisted of a 33% duty cycle (10 seconds on, 20 seconds off) with instantaneous powers of 3, 15, 30, 75 and 150 mW/μm^2^. All embryos were harvested and immediately fixed overnight at 4 °C in 4% formaldehyde in 1x PBS. For the impulse response measurements (Fig. 1 E,F), we injected 15 pg of either Cry-Cib or LOV optoNodal receptors RNAs (i.e. 15 pg each of Type I and Type II receptors) into MZ*oep* embryos at the 1-cell stage. A 20-minute light treatment with average power 20 μW/mm^2^ (60 μW/mm^2^ instantaneous power with 33% duty cycle) was applied beginning at dome stage. At the indicated times, embryos were harvested and fixed overnight at 4 C in 4% formaldehyde in 1x PBS. For both experiments, fixed embryos were immunostained for pSmad2.

### Fixed embryo staining, imaging and quantification

α-pSmad2 immunostaining was performed as previously described^28^. The primary antibody used was CST 18338 at 1:1000 dilution. *Flh*, *lft2, shha*, *sox32* and *myoD* transcripts were detected using an HCR 3.0 protocol^89^. HCR staining was carried out according to manufacturer instructions for <1 dpf zebrafish embryos. Our α-*flh* and α-*lft2* probesets were visualized with AlexaFluor 647-conjugated B3 and AlexaFluor 488-conjugated B2 HCR 3.0 hairpins, respectively. Probes for *shha* and *myoD* were visualized using Alexafluor 546-coupled B2 hairpins, and *sox32* probes were visualized with AlexaFluor 647-coupled B3 hairpins. Both HCR and pSmad2-stained embryos were mounted in 1% low-melt agarose and imaged on Nikon A1 laser scanning confocal microscopes at the University of Pittsburgh Center for Biological Imaging. Z-stacks were acquired with a 2.5 μm spacing on either 10x or 20x air objectives.

α-pSmad2 staining intensity was quantified using a custom MATLAB image analysis pipeline described previously^28^. Briefly, Sytox green-stained nuclei within 25 μm of the embryo animal pole were segmented using a combination of local adaptive thresholding, morphological filtering and active contours boundary refinement. Automated segmentation results were further refined by manual inspection and correction with a custom MATLAB interface. Fluorescence intensities on each imaged channel were compiled for each segmented object. For the quantification panels in Fig. 1, mean pixel intensities within each mask were used. Statistical comparisons between mean intensities of different conditions (e.g. background comparisons in Fig. S1) were performed using an unpaired sample t-test.

### Embryo mount design and fabrication

Molds for embryo ‘egg crate’ mounts were designed using TinkerCad software. Mounts were arrays of short, embryo-sized ‘posts’ with varying radii (275, 300, 325, and 350 μm) and height 600 μm that created individual wells for embryos when molded. Four corner posts of height 3 mm set the spacing between the bottom of the dish and the wells. Each design was exported as toolpath (.stl) files and printed using a Form 3 SLA printer (Formlabs). Some variation in feature dimensions occurs between prints, so the appropriate mold should be selected empirically in a pilot experiment.

To mold embryo eggcrate mounts, 3.0 mL of melted 0.5% agarose in embryo medium was dispensed into a well of a 6-well polystyrene tissue culture plate. The 325 μm egg crate mount mold was placed into the agarose, and excess agarose was removed, to allow the corner spacer legs to contact the bottom of the dish. Mounts were allowed to solidify at 4 °C for > 1 hour, and the mold was manually removed using a scalpel. For patterning experiments, 1-2 mL 0.2% low-melt agarose was layered over the egg crate mount, and embryos were manually loaded into the well array and oriented. The low-melt agarose overlay was allowed to gel for ∼15 minutes at room temperature before mounted embryos were moved to the microscope for optical patterning experiments.

### Patterning endoderm internalization

MZ*oep* embryos were injected with 30 pg of mRNA encoding each optoNodal2 receptor and 150 pg of mCherry mRNA at the 1-cell stage. Embryos were grown in the dark in embryo medium until 3.0 hpf at 28.5 °C, at which point they were transferred into embryo array mounts as described above. Optogenetic treatments were carried out from 3.75-6.25 hpf. Stimulation patterns comprised annular rings at the embryo margin of 75 μm thickness with instantaneous intensity of 240 μW/mm^2^. With scanning over 6 positions, this resulted in an average intensity of 40 μW/mm^2^ (i.e. 16% duty cycle with 20 second dwell time at each position). After stimulation, embryos were immediately retrieved from array mounts and fixed overnight in 4% formaldehyde at 4 °C in the dark. Fixed embryos were stained for *sox32* expression using HCR as described above. Embryos were counterstained with Hoechst nuclear stain. Stained embryos were mounted with the A-V axis parallel to a No. 1.5 glass coverslip in 1% low-melt agarose and imaged on a Nikon A1 confocal with 20x objective on Hoechst and Alexa647 channels. Z-stacks were obtained with 5 μm between slices; presented images are maximum intensity projections.

### Optogenetic rescue of MZ*oep* mutant phenotype

One-cell MZ*oep* embryos were injected with an mRNA cocktail containing mCherry and optoNodal2 receptors. Each embryo received a total of 22.5 pg of each receptor and 150 pg of mCherry mRNAs. Embryos were grown in the dark in agarose-coated 6-well plates containing embryo medium until 3 hpf, at which point they were transferred to embryo array mounts for patterning. Optogenetic stimulation was performed using the mask array depicted in Fig. 4. The average powers of 40, 20 and 10 μW/mm^2^ indicated in the figure were achieved using instantaneous intensities of 240, 120 and 60 μW/mm^2^, respectively (a total of 6 positions were scanned cyclically with a 20 second patterning dwell at each position). To visualize pattern registration, transmitted light (using a 635 nm ‘safe light’ LED positioned under the stage) and fluorescent images (using patterned illumination on the RFP channel) were taken every 15 minutes. After patterning, each embryo was transferred to a well of an agarose-coated 24-well plate containing embryo medium. Transfers were performed to preserve the ordering of embryos in the patterning array; that is, each embryo phenotype could be directly connected back to the live images taken during patterning. Phenotypes were assessed at 26 hpf by mounting embryos laterally in 2.2% methylcellulose and transmitted light imaging on a Leica M165 FC upright microscope. Tissue marker gene expression (e.g. *shha, hgg1,* and *myoD*) was visualized by HCR 3.0 as described above.

### Patterning microscope design

Experiments were performed on two versions of an ultra-widefield patterning microscope. Preliminary experiments were performed on a custom-built design (the ‘Firefly’), described previously^73^. For data shown in this study, we reproduced a Firefly-like microscope using more accessible commercial components. The core of our patterning system was built around a Mightex OASIS Macro DMD microscope. This core system comprised an array of LEDs (405 nm, 470 nm, 560 nm and 625 nm) that were routed to a DMD projector (Mightex Polygon 1000, 1140×912 pixels) via a liquid light guide and a 0.37 NA objective macro lens. The overall magnification of the projection path was 2x, yielding an effective projection ‘pixel size’ of 3.8 μm at the sample plane. The overall imaging of the imaging path of the system is 4x. To facilitate our experiments, we made the following modifications to the system:

### Camera

To facilitate rapid, high-sensitivity imaging, we installed a Hamamatsu Orca Fusion III sCMOS camera in the observation path. The camera was triggered using custom software (see below) via voltage pulses from an Arduino controller through the external trigger port.

### Objective Lens

To control the angular content of incident patterned light, we contracted with Mightex to install a movable iris at the back focal plane of the objective lens. By closing this aperture, the angular content of patterned light could be reduced, resulting in the ‘pencil beam’ configuration used in most patterning experiments in this study. This feature was included in order to render projected patterns less sensitive to the position of an embryo with respect to the objective’s focal plane.

### Filter wheel and main dichroic

To enable multi-channel imaging without channel crosstalk, we installed a large aperture (50 mm) motorized filter wheel (Edmund Optics, 84-889) with DAPI, GFP, RFP and E2-Crimson band emission filters (Chroma). The filter wheel was inserted into the light path using a custom-machined threaded adapter. We replaced the 50-50 beam splitter in the original Oasis Macro design with a large-area, 4-band dichroic (Semrock, DIO3-R405/488/561/635-t3). To minimize pattern distortion due to dichroic curvature over its large area, we selected a 3 mm-thick, 42 x 60 mm material. To fit the dichroic into Macro beam splitter housing, we milled ∼1 mm of excess material out of the Mightex dichroic housing.

### Motorized Stage

To enable automated scanning between multiple positions, we installed a motorized XYZ encoded stage (Prior Instruments, H101E1F XY motor with FB206 focus block stage and ProScan III Controller). Both XY position and Z focal control were managed by moving the sample in 3-dimensions with the stage.

### Sample Incubation

Sample temperature and humidity were controlled during experiments using an OkoLabs BoldLine stage-top incubator system with active humidity and temperature control. Since the microscope has an upright design, patterning and imaging were performed through a transparent lid with active heating to prevent condensation during long experiments.

### Microscope Control

All experiments were performed using a custom-built interface coded in MATLAB. This interface consisted of a custom GUI to streamline (1) DMD calibration, (2) multi-position selection and (3) imaging setting selection. Patterning experiments were performed using custom MATLAB scripts. Our software interface was designed with object classes for the camera, stage, LED array, and DMD instruments. Each object class was designed with high-level class methods to execute hardware commands (e.g. move stage to XYZ position, capture image, activate LED, etc.). Acquisition scripts were built using these high-level methods. The software interface used here was custom-developed in the Lord lab. An open-source software package for control of DMD microscopes (‘Luminos’) has also been recently developed and release by the Cohen lab^90^

### Patterned Illumination

The microscope’s light projection path was calibrated prior to each patterning or acquisition session. To register the DMD projector’s coordinates with spatial coordinates at the sample plane, we projected and imaged a mask containing 10 circular spots with known centroid positions onto a microscope slide with a mirrored surface. This image was then used to fit an affine transformation that maps DMD coordinates to sample plane coordinates. This transform was used to ensure that each projected pattern was properly registered to the targeted spatial coordinates on the sample. To correct an uneven illumination intensity profile, it was measured using static illumination with all pixels ON and imaging its reflection on a mirror. This profile was proportionally applied as the grayscale value of projected masks to achieve a uniform illumination intensity across the FOV. Illumination intensity was measured automatically using a power meter (Thorlabs PM100D), before each experiment at the relevant LED currents.

## Acknowledgements

We thank Travis wheeler at the Universit of Pittsburgh Department of Cell Biology machine shop for support with 3D printing. We also acknowledge the Simon Watkins and the University of Pittsburgh Center for Biological Imaging for support and access to confocal fluorescence imaging resources. This work was supported by a Vannevar Bush Faculty Fellowship grant (N00014-18-1-2859, AEC), an NICHD K99/R00 Award (5K99HD097297, NDL), an NIH R37 Award (GM056211, AFS) and the Chilean National Agency for Research and Development (ANID) Fondo de Desarrollo Cientifico y Tecnológico (Fondecyt 11231198, VJP).

---

**Fig. S1.**
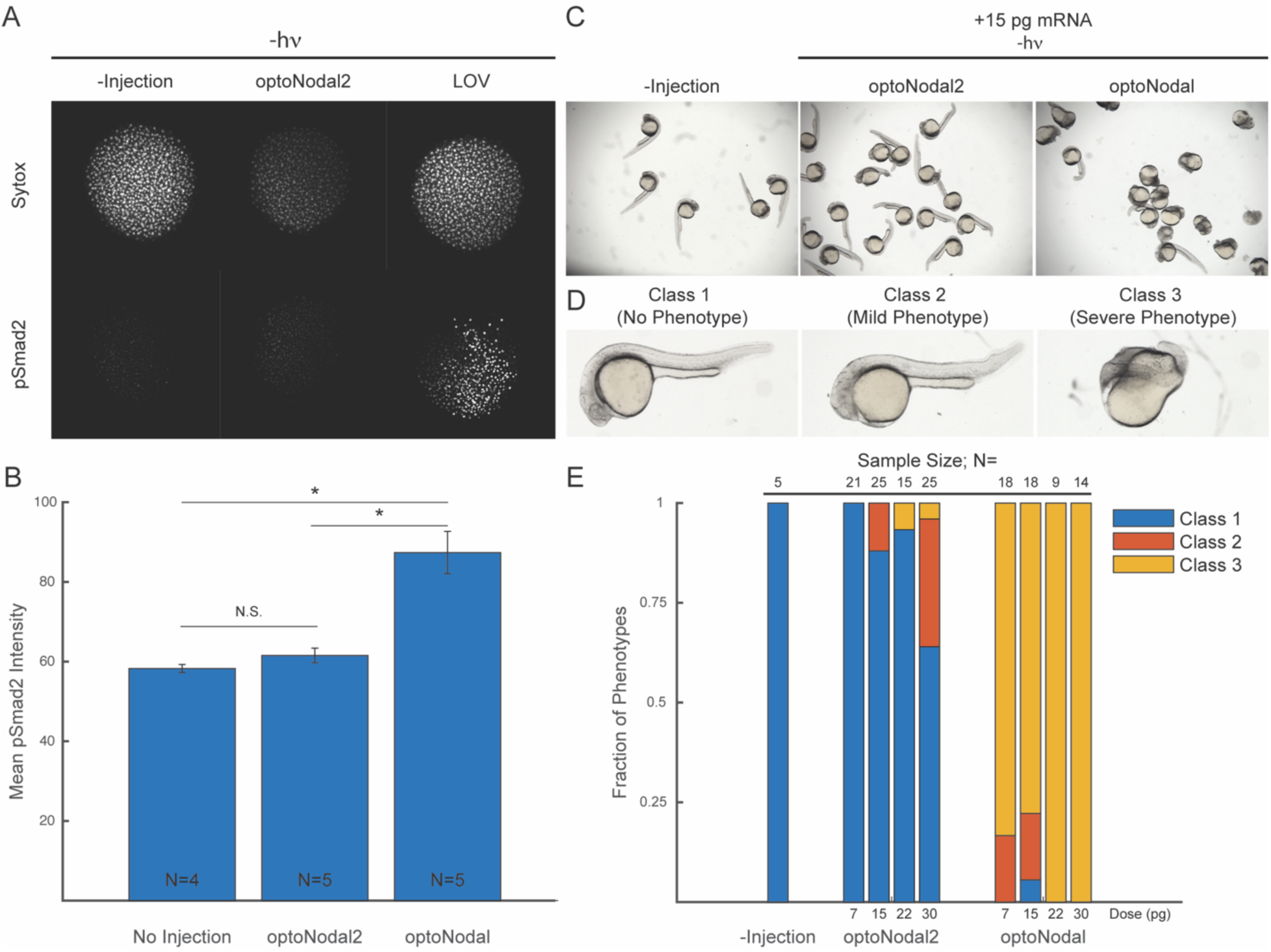
Comparison of dark activity in optoNodal vs. optoNodal2 reagents. (A) Visualization of dark activity. *Mvg1* embryos were unperturbed (‘-Injection’) or injected with mRNA encoding Cry-Cib-based optoNodal2 or LOV-based optoNodal receptors. Embryos were raised in the dark until 5.3 hpf, fixed and immunostained for α-pSmad2 (bottom row). (B) Quantification of α-pSmad2 staining intensity in unilluminated embryos. Graph depicts mean α-pSmad2 nuclear staining intensity, and error bars denote s.e.m.. Statistical comparisons between samples were performed with a unpaired sample t-test with asterisks denoting p < 0.05. (C) Representative 24 hpf phenotypes of wild-type embryos injected with 15 pg of Cry-Cib or LOV-based optoNodal receptors. (D) Example images denoting phenotypic classes quantified in panel E. Class I embryos exhibit no gross abnormalities, Class II embryos exhibit loss of head structures and/or pronounced axis curvature, Class III embryos exhibit severe dorsalization consistent with excess Nodal signaling activity. (E) Distribution of embryos between phenotypic classes in wild-type embryos without injection or injected with indicated amounts of optoNodal2 or optoNodal receptor mRNAs.

**Fig. S2.**
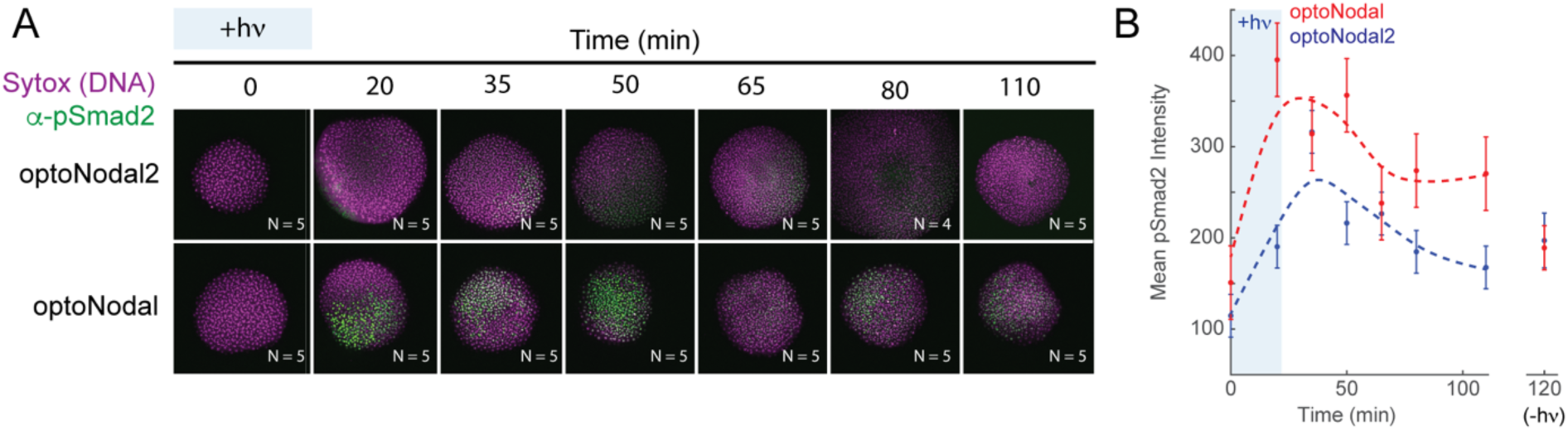
Dynamic responses of optoNodal and optoNodal2 in MZ*oep* mutants. To confirm the observations of Fig. 1 E,F, the responses of optoNodal and optoNodal2 reagents to a 20 minute impulse of light were measured in MZ*oep* mutant embryos. (A) Measurement of response kinetics for optoNodal (top row) and optoNodal2 (bottom row) reagents. Embryos injected with indicated reagents were illuminated for 20 minutes with 470 nm light. Nodal signaling was measured by α-pSmad2 immunostaining (green). Images are maximum intensity projections of representative embryos. (B) Quantification of Nodal signaling activity from panel A. α-pSmad2 staining intensity was extracted from segmented nuclei in optoNodal (red) and optoNodal2 (blue) treatment groups; each point represents the average nuclear staining intensity from the indicated number replicate embryos in Panel A. Error bars denote the standard error of the mean. Background intensity of unilluminated embryos at the 120 minute timepoint are included (-hν) to indicate baseline levels of signaling activity.

**Fig. S3.**
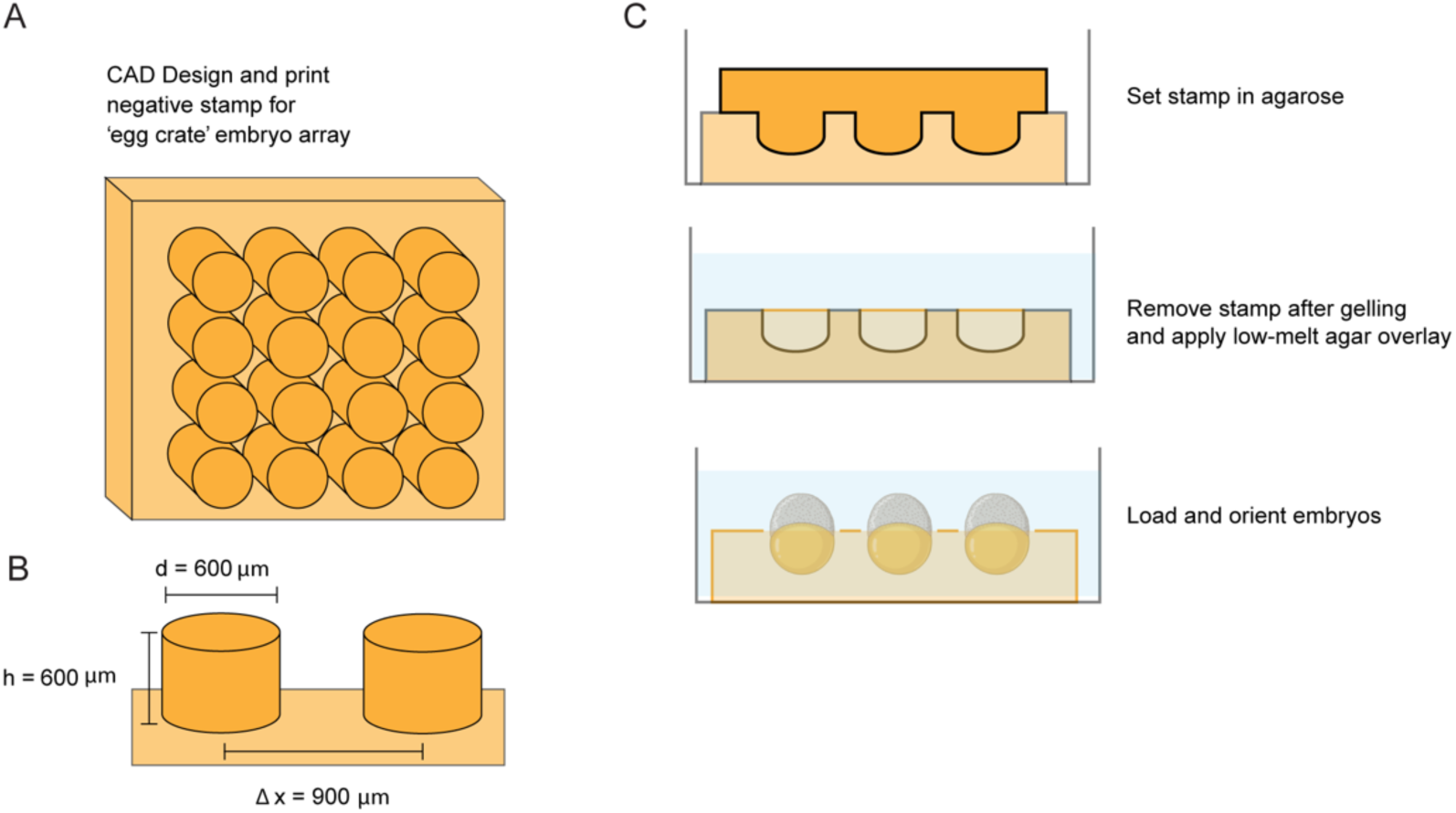
Design and fabrication of embryo array mounts. (A) Design of embryo array mounts. A negative ‘egg crate’ stamp consisting of an array of cylindrical posts was designed using TinkerCAD. (B) Typical dimensions of embryo array stamps. For most experiments, an array of cylindrical posts with 600 μm diameter and height (separated by 300 μm in both dimensions) was used. Stamps were 3D printed using a Form 3 SLA printer. (C) Schematic of procedure used to generate agarose embryo mounts from 3D printed stamps. Stamps were pressed into molten 0.5% agarose in embryo medium. After setting, the stamps were manually removed, and an overlay of 0.2% low-melt agarose in embryo medium was pipetted on top at a temperature of ∼42 °C. Embryos were then mounted in the devices and manually oriented before the low-melt agarose solidified. Once encased between regular and low-melt agarose, embryos were used for patterning experiments.

**Fig. S4.**
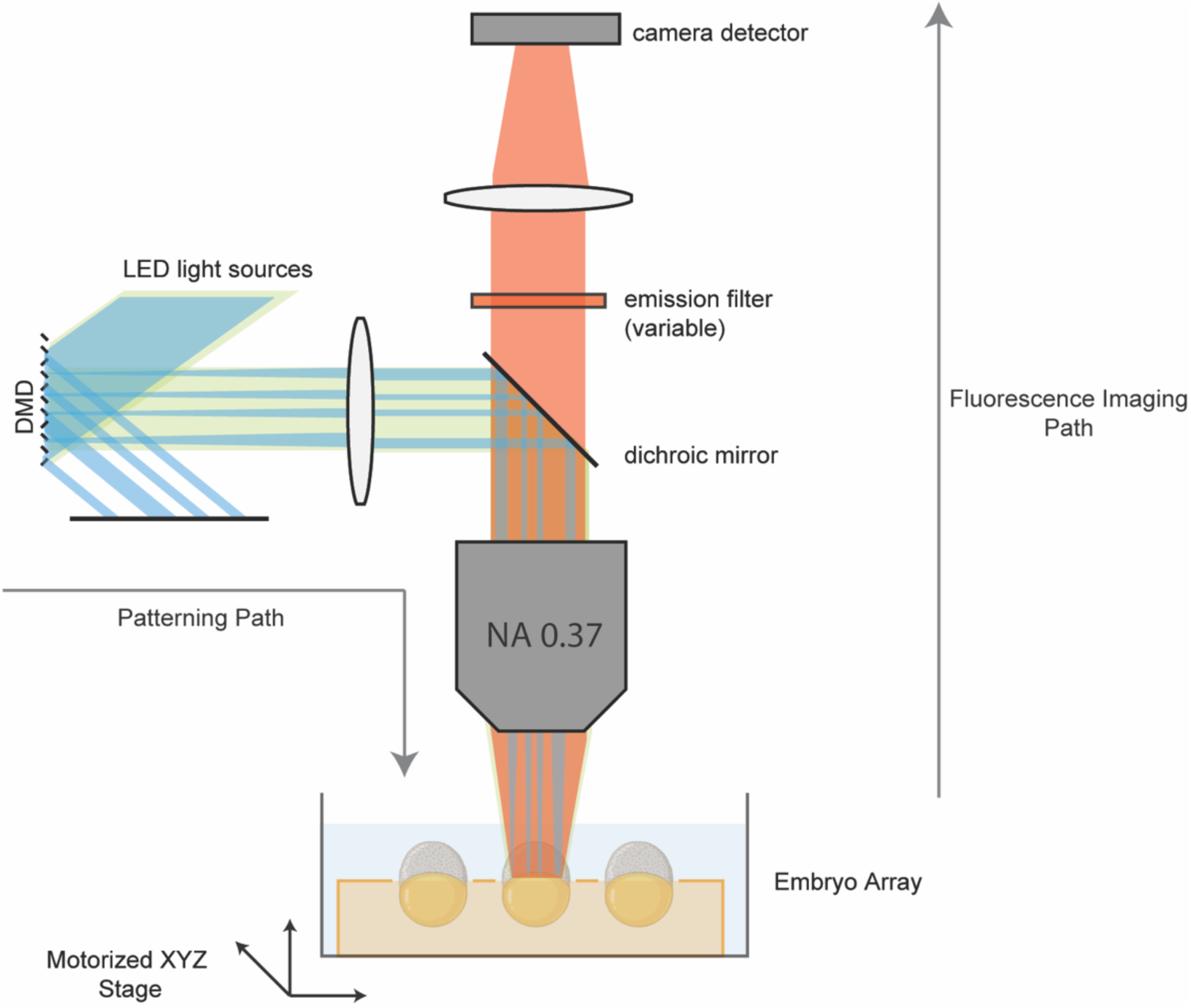
Design of spatial patterning microscope. Our platform is a modified version of the ‘Firefly’ microscope design described by Werley *et al*^91^ and modified by Farhi *et al*^73^. We modified an Oasis Macro ultra-widefield patterning microscope from Mightex. To create spatial patterns at the sample plane (‘Patterning Path’), light from a multi-color LED illuminator is directed to the face of a DMD using a liquid light guide. Pixels on the DMD have two states, ‘ON’ and ‘OFF’, with ‘ON’ pixels directing light toward the sample. Pattern masks are encoded as pixel states on the DMD, and patterned light is collected by a projection lens, reflected off of a multi-band dichroic mirror, and reimaged onto the sample plane using a 4x 0.37 NA objective lens. Emitted light from the sample is collected (‘Fluorescence Imaging Path’) by the objective lens, passed through the dichroic mirror and a wide-format emission filter on a motorized wheel, and reimaged onto a Hamamatsu Orca Fusion III sCMOS camera by a tube lens. Magnification along the projection path (i.e. from DMD to sample plane) is 2x. Magnification along the imaging path (i.e. from sample plane to camera) is 2x. Sample positioning in three dimensions is controlled via an automated XYZ stage.

**<u>Movie S1. Direct visualization of photopatterning of live zebrafish embryos.</u>** To map optical doses delivered to the embryos, wild-type zebrafish embryos were injected at the single-cell stage with mRNA encoding a green-to-red photoconvertible fluorescent protein (Kaede). At sphere stage, embryos were stimulated with 405 nm light with the indicated spatial masks (upper left). Embryos were then imaged every 10 minutes on GFP (upper right) and RFP (lower right) fluorescent channels until 24 hpf.

**<u>Movie S2. Visualization of beating heart tissue in an optogenetically-rescued MZ*oep* mutant.</u>** The 26 hpf embryo was rescued with targeted illumination as described in Fig. 4. Beating heart tissue is located at the midline, suggesting presence of endoderm.

## References

1. Gregor, T., Tank, D. W., Wieschaus, E. F. & Bialek, W. Probing the Limits to Positional Information. Cell 130, 153–164 (2007).

2. Petkova, M. D., Tkačik, G., Bialek, W., Wieschaus, E. F. & Gregor, T. Optimal Decoding of Cellular Identities in a Genetic Network. Cell 176, 844–855.e15 (2019).

3. Zagorski, M. et al. Decoding of position in the developing neural tube from antiparallel morphogen gradients. Science 356, 1379–1383 (2017).

4. Stumpf, H. Mechanism by which cells estimate their location within the body. Nature 212, 430–431 (1966).

5. Wolpert, L. Positional information and the spatial pattern of cellular differentiation. Journal of theoretical biology 25, 1–47 (1969).

6. Driever, W. & Nüsslein-Volhard, C. The *bicoid* protein determines position in the Drosophila embryo in a concentration-dependent manner. Cell 54, 95–104 (1988).

7. Struhl, G., Struhl, K. & Macdonald, P. M. The gradient morphogen *bicoid* is a concentration-dependent transcriptional activator. Cell 57, 1259–1273 (1989).

8. Rogers, K. W. & Schier, A. F. Morphogen gradients: from generation to interpretation. Annual review of cell and developmental biology 27, 377–407 (2011).

9. Kicheva, A. & Briscoe, J. Control of Tissue Development by Morphogens. Annual Review of Cell and Developmental Biology 39, 91–121 (2023).

10. Nemashkalo, A., Ruzo, A., Heemskerk, I. & Warmflash, A. Morphogen and community effects determine cell fates in response to BMP4 signaling in human embryonic stem cells. Development 144, 3042–3053 (2017).

11. Gurdon, J. B. A community effect in animal development. Nature 336, 772–774 (1988).

12. Muncie, J. M. et al. Mechanical tension promotes formation of gastrulation-like nodes and patterns mesoderm specification in human embryonic stem cells. Developmental cell 55, 679–694 (2020).

13. Hagos, E. G. & Dougan, S. T. Time-dependent patterning of the mesoderm and endoderm by Nodal signals in zebrafish. BMC Dev Biol 7, 22 (2007).

14. Johnson, H. E. & Toettcher, J. E. Signaling Dynamics Control Cell Fate in the Early Drosophila Embryo. Developmental Cell 48, 361–370.e3 (2019).

15. Gritsman, K., Talbot, W. S. & Schier, A. F. Nodal signaling patterns the organizer. Development 127, 921–932 (2000).

16. Camacho-Aguilar, E., Yoon, S., Ortiz-Salazar, M. A. & Warmflash, A. Combinatorial interpretation of BMP and WNT allows BMP to act as a morphogen in time but not in concentration. 2022.11.11.516212 Preprint at 10.1101/2022.11.11.516212 (2023).

17. Dessaud, E. et al. Interpretation of the sonic hedgehog morphogen gradient by a temporal adaptation mechanism. Nature 450, 717–720 (2007).

18. Harfe, B. D. et al. Evidence for an Expansion-Based Temporal Shh Gradient in Specifying Vertebrate Digit Identities. Cell 118, 517–528 (2004).

19. Kutejova, E., Briscoe, J. & Kicheva, A. Temporal dynamics of patterning by morphogen gradients. Current Opinion in Genetics & Development 19, 315–322 (2009).

20. Tucker, J. A., Mintzer, K. A. & Mullins, M. C. The BMP Signaling Gradient Patterns Dorsoventral Tissues in a Temporally Progressive Manner along the Anteroposterior Axis. Developmental Cell 14, 108–119 (2008).

21. Sako, K. et al. Optogenetic Control of Nodal Signaling Reveals a Temporal Pattern of Nodal Signaling Regulating Cell Fate Specification during Gastrulation. Cell Reports 16, 866–877 (2016).

22. Heemskerk, I. et al. Rapid changes in morphogen concentration control self-organized patterning in human embryonic stem cells. eLife 8, e40526 (2019).

23. Sorre, B., Warmflash, A., Brivanlou, A. H. & Siggia, E. D. Encoding of temporal signals by the TGF-β pathway and implications for embryonic patterning. Dev Cell 30, 334–342 (2014).

24. Tsai, T. Y.-C. et al. An adhesion code ensures robust pattern formation during tissue morphogenesis. Science 370, 113–116 (2020).

25. Xiong, F. et al. Specified neural progenitors sort to form sharp domains after noisy Shh signaling. Cell 153, 550–561 (2013).

26. Economou, A. D., Guglielmi, L., East, P. & Hill, C. S. Nodal signaling establishes a competency window for stochastic cell fate switching. Developmental cell 57, 2604–2622 (2022).

27. Rogers, K. W. et al. Nodal patterning without Lefty inhibitory feedback is functional but fragile. Elife 6, (2017).

28. Lord, N. D., Carte, A. N., Abitua, P. B. & Schier, A. F. The pattern of nodal morphogen signaling is shaped by co-receptor expression. eLife 10, e54894 (2021).

29. Xu, P.-F., Houssin, N., Ferri-Lagneau, K. F., Thisse, B. & Thisse, C. Construction of a Vertebrate Embryo from Two Opposing Morphogen Gradients. Science 344, 87–89 (2014).

30. Müller, P. et al. Differential diffusivity of Nodal and Lefty underlies a reaction-diffusion patterning system. Science 336, 721–724 (2012).

31. Beyer, H. M. et al. Red Light-Regulated Reversible Nuclear Localization of Proteins in Mammalian Cells and Zebrafish. ACS Synth. Biol. 4, 951–958 (2015).

32. LaBelle, J. et al. TAEL 2.0: An Improved Optogenetic Expression System for Zebrafish. Zebrafish 18, 20–28 (2021).

33. Legnini, I. et al. Spatiotemporal, optogenetic control of gene expression in organoids. Nat Methods 20, 1544–1552 (2023).

34. Rogers, K. W. & Müller, P. Optogenetic approaches to investigate spatiotemporal signaling during development. Current Topics in Developmental Biology 137, 37–77 (2020).

35. Johnson, H. E. & Toettcher, J. E. Illuminating developmental biology with cellular optogenetics. Current opinion in biotechnology 52, 42–48 (2018).

36. Bugaj, L. J., O’Donoghue, G. P. & Lim, W. A. Interrogating cellular perception and decision making with optogenetic tools. Journal of Cell Biology 216, 25–28 (2016).

37. Grusch, M. et al. Spatio-temporally precise activation of engineered receptor tyrosine kinases by light. The EMBO Journal (2014) doi:10.15252/embj.201387695.

38. Kumar, S. & Khammash, M. Platforms for Optogenetic Stimulation and Feedback Control. Front. Bioeng. Biotechnol. 10, (2022).

39. Bugaj, L. J. & Lim, W. A. High-throughput multicolor optogenetics in microwell plates. Nat Protoc 14, 2205–2228 (2019).

40. Repina, N. A. et al. Engineered illumination devices for optogenetic control of cellular signaling dynamics. Cell reports 31, (2020).

41. Johnson, H. E. et al. The Spatiotemporal Limits of Developmental Erk Signaling. Developmental Cell 40, 185–192 (2017).

42. Johnson, H. E., Djabrayan, N. J. V., Shvartsman, S. Y. & Toettcher, J. E. Optogenetic Rescue of a Patterning Mutant. Current Biology 30, 3414–3424.e3 (2020).

43. Ho, E. K. et al. Dynamics of an incoherent feedforward loop drive ERK-dependent pattern formation in the early Drosophila embryo. Development 150, dev201818 (2023).

44. Bugaj, L. J., Choksi, A. T., Mesuda, C. K., Kane, R. S. & Schaffer, D. V. Optogenetic protein clustering and signaling activation in mammalian cells. Nat Methods 10, 249–252 (2013).

45. Singh, A. P. et al. Optogenetic control of the Bicoid morphogen reveals fast and slow modes of gap gene regulation. Cell Reports 38, 110543 (2022).

46. Humphreys, P. A. et al. Optogenetic Control of the BMP Signaling Pathway. ACS Synth. Biol. 9, 3067– 3078 (2020).

47. Čapek, D. et al. Light-activated Frizzled7 reveals a permissive role of non-canonical wnt signaling in mesendoderm cell migration. Elife 8, e42093 (2019).

48. Rogers, K. W., ElGamacy, M., Jordan, B. M. & Müller, P. Optogenetic investigation of BMP target gene expression diversity. eLife 9, e58641 (2020).

49. Schier, A. F. Nodal signaling in vertebrate development. Annual review of cell and developmental biology 19, 589–621 (2003).

50. Chen, Y. & Schier, A. F. The zebrafish Nodal signal Squint functions as a morphogen. Nature 411, 607– 610 (2001).

51. Conlon, F. L. et al. A primary requirement for nodal in the formation and maintenance of the primitive streak in the mouse. Development 120, 1919–1928 (1994).

52. Feldman, B. et al. Zebrafish organizer development and germ-layer formation require nodal-related signals. Nature 395, 181–185 (1998).

53. Gritsman, K. et al. The EGF-CFC protein one-eyed pinhead is essential for nodal signaling. Cell 97, 121–132 (1999).

54. Reissmann, E. et al. The orphan receptor ALK7 and the Activin receptor ALK4 mediate signaling by Nodal proteins during vertebrate development. Genes Dev. 15, 2010–2022 (2001).

55. Yeo, C.-Y. & Whitman, M. Nodal Signals to Smads through Cripto-Dependent and Cripto-Independent Mechanisms. Molecular Cell 7, 949–957 (2001).

56. Attisano, L. & Wrana, J. L. Signal Transduction by the TGF-β Superfamily. Science 296, 1646–1647 (2002).

57. Massagué, J., Seoane, J. & Wotton, D. Smad transcription factors. Genes & development 19, 2783– 2810 (2005).

58. Dubrulle, J. et al. Response to Nodal morphogen gradient is determined by the kinetics of target gene induction. eLife 4, e05042 (2015).

59. Dougan, S. T., Warga, R. M., Kane, D. A., Schier, A. F. & Talbot, W. S. The role of the zebrafish nodal-related genes squint and cyclops in patterning of mesendoderm. (2003).

60. Erter, C. E., Solnica-Krezel, L. & Wright, C. V. Zebrafish nodal-related 2Encodes an early mesendodermal inducer signaling from the extraembryonic yolk syncytial layer. Developmental biology 204, 361–372 (1998).

61. Rebagliati, M. R., Toyama, R., HaÑer, P. & Dawid, I. B. Cyclops encodes a nodal-related factor involved in midline signaling. Proceedings of the National Academy of Sciences 95, 9932–9937 (1998).

62. Sampath, K. et al. Induction of the zebrafish ventral brain and floorplate requires cyclops/nodal signalling. Nature 395, 185–189 (1998).

63. Montague, T. G. & Schier, A. F. Vg1-Nodal heterodimers are the endogenous inducers of mesendoderm. Elife 6, e28183 (2017).

64. Pelliccia, J. L., Jindal, G. A. & Burdine, R. D. Gdf3 is required for robust Nodal signaling during germ layer formation and left-right patterning. Elife 6, e28635 (2017).

65. Bisgrove, B. W., Su, Y.-C. & Yost, H. J. Maternal Gdf3 is an obligatory cofactor in Nodal signaling for embryonic axis formation in zebrafish. eLife 6, e28534 (2017).

66. Thisse, B., Wright, C. V. & Thisse, C. Activin-and Nodal-related factors control antero–posterior patterning of the zebrafish embryo. Nature 403, 425–428 (2000).

67. Vincent, S. D., Dunn, N. R., Hayashi, S., Norris, D. P. & Robertson, E. J. Cell fate decisions within the mouse organizer are governed by graded Nodal signals. Genes & development 17, 1646–1662 (2003).

68. Schier, A. F., Neuhauss, S. C. F., Helde, K. A., Talbot, W. S. & Driever, W. The one-eyed pinhead gene functions in mesoderm and endoderm formation in zebrafish and interacts with no tail. Development 124, 327–342 (1997).

69. Pinheiro, D., Kardos, R., Hannezo, É. & Heisenberg, C.-P. Morphogen gradient orchestrates pattern-preserving tissue morphogenesis via motility-driven unjamming. Nature Physics 18, 1482–1493 (2022).

70. Carmany-Rampey, A. & Schier, A. F. Single-cell internalization during zebrafish gastrulation. Current biology 11, 1261–1265 (2001).

71. Takahashi, F. et al. AUREOCHROME, a photoreceptor required for photomorphogenesis in stramenopiles. Proceedings of the National Academy of Sciences 104, 19625–19630 (2007).

72. Pudasaini, A., El-Arab, K. K. & Zoltowski, B. D. LOV-based optogenetic devices: light-driven modules to impart photoregulated control of cellular signaling. Front Mol Biosci 2, 18 (2015).

73. Farhi, S. L. et al. Wide-area all-optical neurophysiology in acute brain slices. Journal of Neuroscience 39, 4889–4908 (2019).

74. Li, Y. et al. Spatiotemporal Control of TGF-β Signaling with Light. ACS Synth. Biol. 7, 443–451 (2018).

75. Kennedy, M. J. et al. Rapid blue-light–mediated induction of protein interactions in living cells. Nature methods 7, 973–975 (2010).

76. Jia, B. Z., Qi, Y., David Wong-Campos, J., Megason, S. G. & Cohen, A. E. A bioelectrical phase transition patterns the first beats of a vertebrate heart. doi:10.1101/2022.12.06.519309.

77. Liu, Z., Woo, S. & Weiner, O. D. Nodal signaling has dual roles in fate specification and directed migration during germ layer segregation in zebrafish. Development 145, dev163535 (2018).

78. Emig, A. A. et al. Temporal dynamics of BMP/Nodal ratio drive tissue-specific gastrulation morphogenesis. 2024.02.06.579243 Preprint at 10.1101/2024.02.06.579243 (2024).

79. van Boxtel, A. L., Economou, A. D., Heliot, C. & Hill, C. S. Long-Range Signaling Activation and Local Inhibition Separate the Mesoderm and Endoderm Lineages. Developmental Cell 44, 179–191.e5 (2018).

80. Crick, F. Diffusion in Embryogenesis. Nature 225, 420–422 (1970).

81. Wartlick, O., Kicheva, A. & González-Gaitán, M. Morphogen Gradient Formation. Cold Spring Harb Perspect Biol 1, a001255 (2009).

82. Yu, S. R. et al. Fgf8 morphogen gradient forms by a source-sink mechanism with freely diffusing molecules. Nature 461, 533–536 (2009).

83. Kerszberg, M. & Wolpert, L. Mechanisms for Positional Signalling by Morphogen Transport: a Theoretical Study. Journal of Theoretical Biology 191, 103–114 (1998).

84. Müller, P., Rogers, K. W., Yu, S. R., Brand, M. & Schier, A. F. Morphogen transport. Development 140, 1621–1638 (2013).

85. Ochoa-Espinosa, A., Yu, D., Tsirigos, A., Struffi, P. & Small, S. Anterior-posterior positional information in the absence of a strong Bicoid gradient. Proceedings of the National Academy of Sciences 106, 3823–3828 (2009).

86. The Zebrafish Book; A guide for the laboratory use of zebrafish (Danio rerio) | CiNii Research. https://cir.nii.ac.jp/crid/1370283694361132063.

87. Kimmel, C. B., Ballard, W. W., Kimmel, S. R., Ullmann, B. & Schilling, T. F. Stages of embryonic development of the zebrafish. Developmental Dynamics 203, 253–310 (1995).

88. Thomas, O. S., Hörner, M. & Weber, W. A graphical user interface to design high-throughput optogenetic experiments with the optoPlate-96. Nat Protoc 15, 2785–2787 (2020).

89. Choi, H. M. T. et al. Third-generation in situ hybridization chain reaction: multiplexed, quantitative, sensitive, versatile, robust. Development 145, dev165753 (2018).

90. Luminos: bi-directional microscopy software. https://www.luminosmicroscopy.com/

91. Werley, C. A., Chien, M.-P. & Cohen, A. E. Ultrawidefield microscope for high-speed fluorescence imaging and targeted optogenetic stimulation. Biomedical optics express 8, 5794–5813 (2017).

